# A thermal time framework drives coordinated below- and above-ground development in temperate cereal crops

**DOI:** 10.64898/2026.04.24.720680

**Authors:** Matías Schierenbeck, Akshay B. Tawale, Ivan Lopez-Valdivia, Dylan H. Jones, Annegret Wolf, Petra Linow, Corinna Trautewig, Hannah M. Schneider

**Author notes:** Authors for correspondence: Matías Schierenbeck and Hannah M. Schneider.

## Abstract

- Cereal architecture is underpinned by the coordinated development of modular phytomer units. While above-ground phenology is well characterized by metrics such as the phyllochron, an equivalent framework for root system development is lacking. Because each phytomer node initiates both leaves and adventitious roots, root and shoot development are inherently linked.
- Here, we quantified this coordination in wheat, barley, and rye across contrasting temperature regimes and validated the results under field conditions. We introduce the rhizochron, defined as the thermal time (growing degree-days, °C d) period between the emergence of nodal roots on successive stem nodes, and the root appearance interval, describing the emergence rate of individual root axes. Root development followed a highly conserved thermal sequence synchronized with shoot phenology.
- Across species and environments, the rhizochron averaged 146.1°C d, closely matching the phyllochron (126.6°C d). We also identified a consistent thermal offset, with nodal roots emerging approximately 185.3°C d after the corresponding leaf on the same phytomer node. The root appearance interval averaged 45.3°C d, reflecting continuous root deployment across active nodes.
- By integrating root phenology into a node-based framework, the rhizochron provides a predictive tool for crop modeling, trait-based breeding, and more target phenotyping aimed at improving resource acquisition and climate resilience.

## Introduction

A fundamental feature of plant development is the coordinated production of organs and tissues across the entire organism. Cereal plant architecture is fundamentally defined by the integration of modular structural units and the thermal time governing their appearance. The phytomer serves as the basic building block of this assembly, representing a repeating unit consisting of a node, an internode, a leaf, an axillary bud, and nodal (i.e. adventitious) roots (Wilhelm and McMaster, 1995; Forster et al, 2007). The systematic accumulation of these units constructs the physical framework of the shoot, with their rate of appearance dictated by the phyllochron, the thermal time interval, typically expressed in growing degree-days (GDD), between the emergence of successive leaves (Kirby et al. 1985). Because the leaf is the most readily observable component of the phytomer, the phyllochron has historically functioned as the primary timing mechanism for modeling shoot development. This thermal synchronization ensures that the addition of new nodes, and the potential tillers and leaves associated with them, remains coupled with environmental signals, providing a predictable metric for shoot phenology across varying climates (Roth et al. 2024).

Leaf appearance follows a relatively stable phyllochron of approximately 60 to 120 °C d per leaf depending on genotype, growth habit, sowing date, and environmental conditions (Kirby et al., 1985; Jamieson et al. 1995; Slafer & Rawson, 1997). This above-ground phenology can be precisely categorized by scales such as Zadoks (Zadoks et al., 1974), the BBCH (Biologische Bundesanstalt, Bundessortenamt und CHemische Industrie) scale (Lancashire et al., 1991), and the Feekes scale (Large, 1954). Together, these approaches enable a highly predictable and transferable description of shoot development across environments.

While the phyllochron provides a visible proxy for vegetative development, the underlying coordination of phytomer formation and stem development is less readily quantified. Recent evidence in barley (*Hordeum vulgare* L.) indicates that node initiation and internode elongation are tightly coordinated processes that together define shoot architecture (Huang et al., 2024). This highlights that the phyllochron reflects not only leaf appearance, but broader developmental dynamics within the phytomer. Accordingly, plant architecture can be viewed not simply as a static form, but as a dynamic system of coordinated phytomeric units (Huang et al., 2024).

Despite this established framework for above-ground development, a similar metric for the thermal-time-driven emergence of axial roots has remained largely undefined. Roots play a fundamental role in the productivity and resilience of cereal crops, such as wheat (*Triticum aestivum* L.), barley, and rye (*Secale cereale* L.) (Jones et al. 2026). Beyond anchoring the plant, root systems regulate water and nutrient uptake, contribute to soil structure formation, interact with the microbiome, and play a pivotal role in carbon sequestration (Gregory et al., 2009; Bardgett et al., 2014). These functions directly influence crop performance and resilience in increasingly variable climates (Schneider & Lynch, 2020). Efficient root systems are of particular importance for the acquisition of nutrients such as nitrogen and phosphorus, and deep root systems enhance tolerance to water deficit by enabling access to subsoil moisture during drought (Lynch, 2019).

The cereal root system is a fibrous root system that initiates with one primary and subsequently typically many seminal roots emerging directly from the seed. As the plant develops, it produces adventitious (i.e. nodal) roots from below-ground stem nodes of the main stem and tillers (Jones et al. 2025, 2026) (Figure 1). Each phytomer node also serves as the initiation point for these nodal roots, suggesting that root system development is intrinsically linked to the same modular growth pattern as the shoot (Nemoto et al. 1995; Yang et al. 1998; Gonin et al. 2019). Comparative studies between the grass genetic model *Brachypodium distachyon* and wheat provide foundational evidence that this coordination is highly conserved; in both species, leaves and nodal roots emerge in a synchronized manner across diverse environments. Specifically, nodal roots develop at specific leaf stages, with nodal roots (in this case coleoptile node roots) appearing at the third leaf stage and multiple nodal roots emerging from multiple stem nodes following the appearance of the fourth leaf (Watt et al 2009). Singh et al. (2010) demonstrated that the specific timing of these events varies significantly between species, with maize (*Zea mays* L.) initiating nodal roots at the 2^nd^ leaf stage while in sorghum (*Sorghum bicolor* L.) this appearance is delayed until the 4^th^ to 5^th^ leaf stage. Similar approaches have been reported for rice (*Oryza sativa*) (Nemoto et al. 1995) and forage grasses, including perennial ryegrass (*Lolium perenne*) and tall fescue (*Festuca arundinacea*) (Yang et al. 1998; Robin et al. 2021)

**Figure 1.**
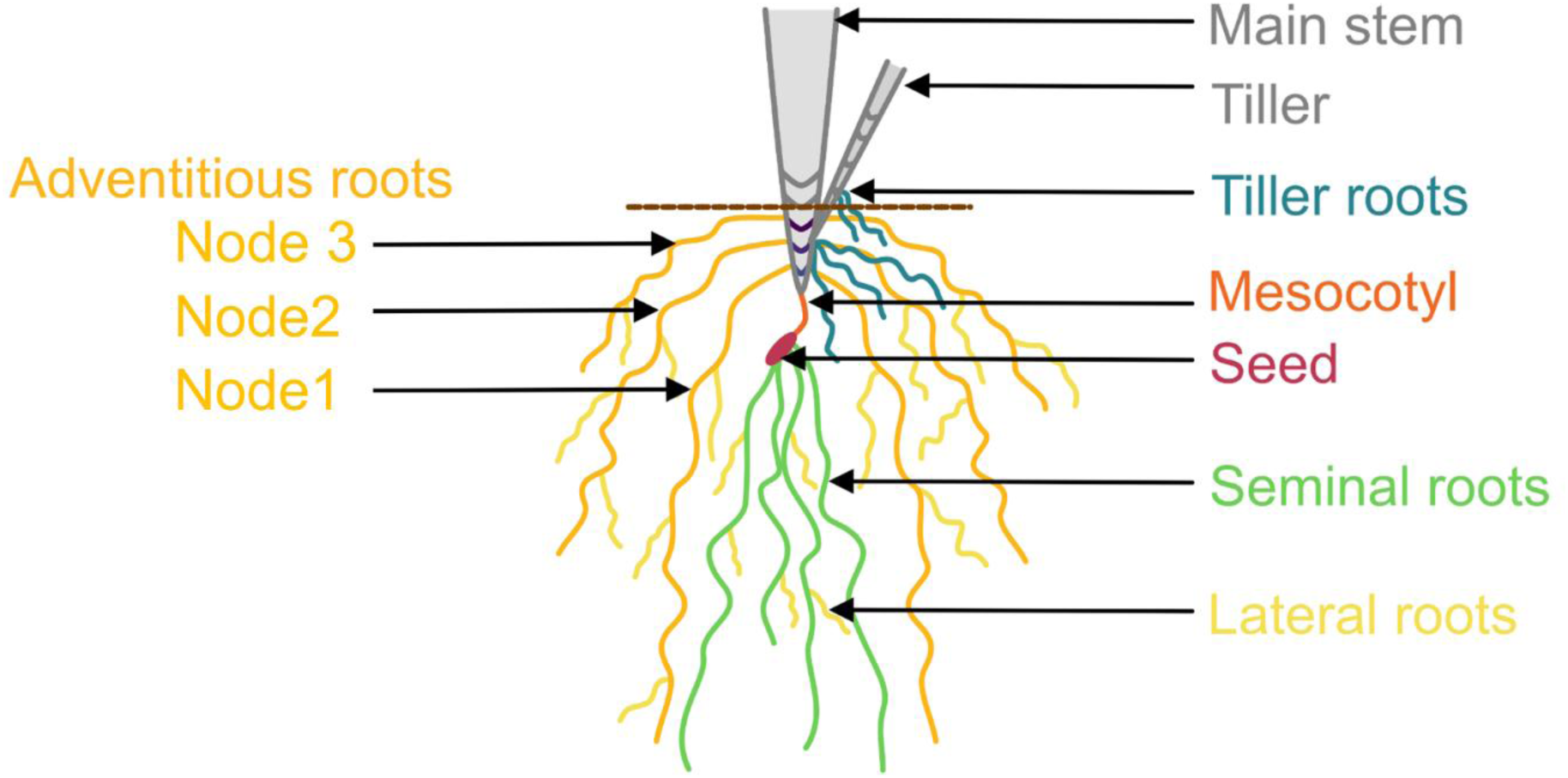
Monocot root system. The figure depicts the fibrous root system architecture of tillering temperate cereals. Seminal roots (seed-borne), adventitious (nodal) roots (stem-borne), and lateral roots (root-borne) are labeled. Adapted from Jones et al., 2026.

While these observations suggest a precise phenological alignment between organ systems, Vincent and Gregory (1989 a, b) established that wheat leaf and nodal root axis production are linearly coordinated by thermal time (base temperature = 0 °C), regardless of nitrogen levels or sowing date. However, while temperature dictates the modular timing of root initiation, the ultimate extension (i.e. growth of a root axis after emergence) and dry matter accumulation of the system are regulated by assimilate supply and show a linear relationship with intercepted photosynthetically active radiation (PAR) (Vincent and Gregory, 1989a, b). While these initial observations suggest a precise temporal alignment between above- and below-ground organ system development, the quantitative details regarding the exact thermal intervals and hierarchical emergence patterns of roots across successive individual nodes as the plant scales its architecture remain to be fully characterized.

Early modeling approaches, such as *WHTROOT,* assumed a degree of synchronization between shoot and root development by parameterizing root growth as a function of thermal time (Porter et al., 1986; Klepper et al., 1997). However, most subsequent research and modeling frameworks shifted toward representing root development primarily as a response to resource availability or biomass allocation, rather than as an explicit developmental process (Asseng et al. 1997; Dupuy et al. 2010). Consequently, the emergence of roots is rarely described as a thermal-time-driven event. While experimental studies in temperate cereals suggest that nodal root formation declines during the transition from vegetative to reproductive growth (Gregory et al. 1978; Barraclough, 1984), empirical quantification of this coordination across species and temperature regimes remains limited.

We investigated whether root development can be systematically linked to thermal time and synchronized with shoot phenology. To facilitate this, we utilize the rhizochron as a framework for measuring the thermal time interval between the emergence of roots on successive stem nodes (cf. Robin et al 2021). Analogous to the phyllochron’s tracking of leaf appearance, the rhizochron quantifies when roots emerge from successive nodes. Establishing this metric enables a whole-plant phenological perspective, mathematically relating root emergence at specific nodes to leaf development. Through sequential harvests of wheat, barley, and rye, we evaluated whether nodal root emergence follows predictable trajectories across temperature regimes to provide a quantitative bridge between below-ground phenology and established shoot development stages.

## Materials and methods

### Experimental site and growth conditions

This study was conducted in controlled-environment greenhouses at the Leibniz Institute of Plant Genetics and Crop Plant Research (Gatersleben, Germany). To evaluate the impact of thermal time accumulation rates on plant development, two distinct temperature regimes were established under a 16-hour photoperiod. The cool chamber was maintained at 19/15 °C (day/night; average 18 °C over 24-h) with 850-1000 µmol m^-2^s^-1^ light intensity and 70% relative humidity, yielding a daily accumulation of 18 growing degree-days (GDD), while the warm chamber was set to 28/22 °C (day/night; average 26 °C over 24-h), resulting in 26 GDD per day. Thermal time was expressed as accumulated GDD, calculated from the daily mean air temperature (T) as GDD= T_mean−T_base (0 °C) (Porter and Gawith, 1999).

### Experimental design and plant material

The experiment followed a split-split plot design with three replications (n=432). Temperature regimes served as main plots, crop species (spring wheat, two-row spring barley, and spring rye) as subplots, and genotypes as sub-subplots. For each species, three genotypes were selected. Wheat: *KWS Sharki* (Germany), *Baguette 801 Premium* (Argentina) and *Klein Vencedor* (Argentina); Barley: *RGT Planet* (Germany), *Quilmes Carisma* (Argentina) and *Isaria* (Germany) Rye: *Sorom* (Germany), *Petka* (Germany) and *Petkuser* (Germany). Seeds were provided by the Federal Ex-Situ German Genebank (IPK Gatersleben).

Sampling occurred at eight sequential intervals every 180 GDD, spanning from the one-expanded-leaf stage (Zadoks stage Z11) through anthesis (Zadoks stage Z65). These sampling times covered major phenological phases: seedling and tillering (180, 360, 540 GDD), stem elongation to flag leaf expansion (720, 900 GDD), booting (1080, 1260 GDD), and heading to anthesis (1440 GDD). For the 18 °C chamber, the experiment duration was 80 days. For the 26 °C chamber, the experiment duration was 55 days. Throughout the manuscript, growing degree days are expressed as thermal time accumulated from the day of transplanting in the solution culture.

### Solution culture

Seeds of each genotype were sterilized (5 minutes in 10% NaClO) and placed in tap water humidified filter paper. Seedlings were germinated in a growth chamber at 22 °C, 60% relative humidity, and 16 h light/8 h dark at a light density of 400 µmol m^-2^s^-1^ (ISTA protocol, 2013). After one week (94 GDD after germination), seedlings were transplanted in the greenhouse (conditions described above) into 16, 75.5 L solution culture systems (Rubbermaid Brute® containers; 38.3 × 70.7 × 44.1 cm) containing nutrient solution (6 mM NH_4_NO_3_, 4 mM MgSO_4_, 2 mM KH_2_PO_4_, 0.3mM Fe(III)-DTPA, 46 μmol H_3_BO_3_, 9 μmol MnCl_2_ 4H_2_O, 0.77 μmol ZnSO_4_ 7H_2_O, 0.32 μmol CuSO_4_ 5H_2_O, 0.07 μmol SO_4_ 5H_2_O; pH 6.4) and equipped with a continuous aeration system. Each container lid was fitted with fifteen evenly spaced planting sites in a 5-by-3 grid pattern, each with a diameter of 4.5 cm. The plants were supported by specially designed 3D printed plastic holders (Supplementary Figure 1).

### Shoot and root measurements

At each sampling point (i.e. every 180 GDD) above- and below-ground development and growth was measured. Above-ground development was scored using the Zadoks scale (Zadoks et al. 1974). The main stem was defined as the primary culm that originated directly from the seed embryo. At each sampling point, the number of leaves on the main stem was counted and the phyllochron was calculated (described below). Total biomass of the above-ground main stem (oven-dried at 60 °C for 72 hours) was collected.

We recorded three types of root data per sampling time: (1) number of seminal roots (S), (2) number of nodal roots per individual node on the main stem (N1 through N6) (i.e. number of roots per node), and (3) total nodal roots from tillers. Roots were considered emerged when longer than 2 cm. The number of main stem nodes with emerged roots was also recorded. All roots from each subclass (e.g. seminal roots, roots from main stem nodes 1, 2, 3, 4, 5, and 6, and nodal roots from tillers) were scanned on a flatbed scanner (Epson Expression 12000XL) at 600 DPI and architectural traits, including total root surface area, were quantified using RhizoVision Explorer 2.0.3 (Seethepalli et al. 2021). Following imaging, all individual root categories were separately oven-dried at 60 °C for 72 hours to determine dry weight. Root and shoot development and growth measurements (Supplementary Table S1) were parameterized in *OpenSimRoot* (Postma et al, 2017), a functional-structural plant soil model to create a visual graphic of temperate cereal root development and growth to supplement the Zadocks scale. The model used a customized version of the input generation scripts described in Lopez-Valdivia et al., (2025, 2023)

### Root development scale

We used a root scale notation (e.g., N5.4) to define developmental stages. The first value indicates the number of main stem nodes with emerged roots; the second value denotes the number of roots on the youngest node where root emergence has occurred. In the example N5.4, the plant has five root-bearing nodes, with four roots present on the fifth node.

### Calculation of the Phyllochron, Rhizochron, and Root Appearance Interval

To maintain developmental consistency across species and treatments, three thermal metrics were calculated using cumulative GDD. The phyllochron is defined as the thermal interval between the appearance of successive leaves on the main stem (Kirby et al. 1985) (Equation 1) and was calculated between 180 and 900 GDD. This specific shortened phenological window was targeted because main stem leaf development ceases after approximately 900 GDD once the final leaf (flag leaf) emerges (*ca.* flag leaf stage, Z39).

The appearance rates for leaves, nodal root positions, and total roots were determined by fitting linear regression models to the relationship between organ count (N) and cumulative thermal time (GDD). From these models, the phyllochron, rhizochron, and root appearance interval were derived as the mathematical inverse of the regression slopes (Equations 1–3), representing the thermal time required for the appearance of successive units (*d*GDD/*d*N; cf. Hay & Kirby, 1991). While the phyllochron was calculated for early vegetative growth, the rhizochron and root appearance interval were evaluated over a broader window (180–1440 GDD) to capture the protracted nature of below-ground development.

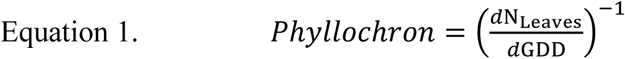

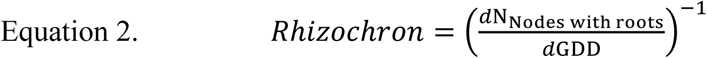

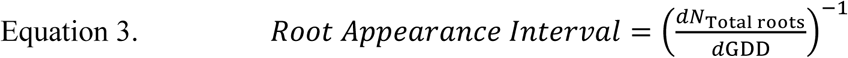

The coordination ratio was calculated as the ratio of rhizochron to phyllochron, with both parameters expressed in thermal time (°C d; GDD) from 180 to 900 GDD. This ratio quantifies the relative timing of root and shoot organ emergence and was computed per plant and averaged across replicates, species, and genotypes.

### Field experiments: validation of the root appearance interval and rhizochron

To validate the rhizochron and root appearance interval, a field experiment was carried out at the Leibniz Institute of Plant Genetics and Crop Plant Research (IPK) (Gatersleben, Germany) (11° 16’ LE; 51°49′ LN) during 2025. The spring wheat variety, Quintus (Saaten Union GmbH, Germany), was sown in 1.2 m^2^ plots (1 m long by 1.2 m wide), containing six rows and a seeding rate of 300 seeds m^-2^. The crop was managed according to local agronomic practices. These standard practices comprise a nitrogen fertilization scheme: 60 kg ha^-1^ at the beginning of the vegetative period (Z13, Zadoks et al. 1974), 60 kg ha^-1^ during stem elongation (Z31-32) and 60 kg ha^-1^ during booting stage (GS47-49). Standard agronomic practices for managing insects and weeds were implemented throughout the crop cycle. Plots were sprayed against weeds with 5 g a.i. ha^−1^ florasulam + 45 g a.i. ha^−1^ pinoxaden (Axial komplett®); a mixture of carfentrazone-ethyl (20 g a.i. ha⁻¹; Aurora 40 WG) and mecoprop-P (700 g a.i. ha⁻¹) + 2,4-D (320 g a.i. ha⁻¹; Duplosan KV neu). For pest management, deltamethrin (7.5 g a.i. ha⁻¹; Decis Forte®) was applied.

Weekly plant evaluation and root crown excavations (Trachsel et al. 2011) were conducted to assess above- and below-ground phenological development across eleven sampling points (spanning 72 days) covering crop emergence to the anthesis period (42 to 1082 GDD, calculated from emergence). The number of emerged roots (> 2 cm in length) on each stem node, the number of stem nodes with emerged roots, and the number of leaves on the main stem were recorded at each sample point. Daily minimum, maximum and mean air temperature (°C) were recorded from nearby meteorological stations. Cumulative GDD was calculated taking into consideration the mean daily air temperature.

### Calculation of temporal offset between leaf and root emergence at individual phytomer nodes

The temporal offset between leaf and root emergence on the same phytomer was determined using linear regression analysis based on accumulated GDD. For each genotype, the number of nodes with emerged leaves and roots across three biological replicates were averaged at each sampling point defined by accumulated GDD. To ensure the accuracy of the calculated emergence rates, we restricted the regression to the active, linear growth phase (180-540 GDD). Ordinary least squares regression was applied to model the number of emerged nodes (N) with leaves (L) and with roots (R) as a function of GDD for both leaves (N_L_ = m_L_ * GDD + b_L_) and roots (N_R_ = m_R_ * GDD + b_R_), yielding organ-specific emergence rates (slopes, m) and initial developmental delays (y-intercepts, b). The exact thermal time of emergence for any specific target node (k) was derived by rearranging the respective regression equations to solve for GDD; specifically, GDD_L,k_ = (k – b_L_) / m_L_) for leaves and GDD_R,k_ = (k – b_R_) / m_R_) for roots. Finally, the temporal offset (ΔGDD) at target node k was calculated as the difference between the thermal time of root emergence and leaf emergence (ΔGDD_k_ = GDD_R,k_ - GDD_L,k_). Under this framework, a positive ΔGDD indicates that the leaf emerged prior to the root on that specific phytomer.

All statistical analyses and graphs were prepared in R (v4.5.3) (R Core Team, 2026) using the tidyverse (v2.0.0) suite (Wickham et al. 2016, 2019) (ggplot2 v4.0.2, dplyr v1.2.1, readr v2.2.0, tidyr v1.3.2) and ggpmisc (v0.7.0) (Aphalo, 2026) on Rstudio (Posit Team, 2026). Two–way ANOVA models were fitted as linear models and post–hoc pairwise comparisons of estimated marginal means were performed with emmeans (v2.0.2) (Lenth & Piaskowski, 2026) using the Sidak adjustment. Model assumptions (independence, approximate normality of residuals, and homogeneity of variances) were checked by residual diagnostics.

## Results

### Relative rates of organ appearance

The developmental coordination between shoot and root emergence in this study showed varying degrees of sensitivity to species and thermal regimes. The phyllochron exhibited a significant species × temperature interaction (p ≤ 0.001), with distinct responses across barley, rye, and wheat. No significant differences were detected between temperature regimes (p = 0.578) (Figure 2, Table 1, Supplementary Table S1, S3). In the greenhouse environment, the mean phyllochron was approximately 126.6 °C d leaf^-1^. In contrast, the rhizochron (the thermal interval between the emergence of roots on successive stem nodes) averaged 146.1 °C d node^-1^; however, it was not significantly influenced by species (p = 0.674), species × temperature interaction (p = 0.800) or temperature (p = 0.839) within this experiment (Figure 2, Supplementary Figure 3, Supplementary Table S1, S3).

**Figure 2.**
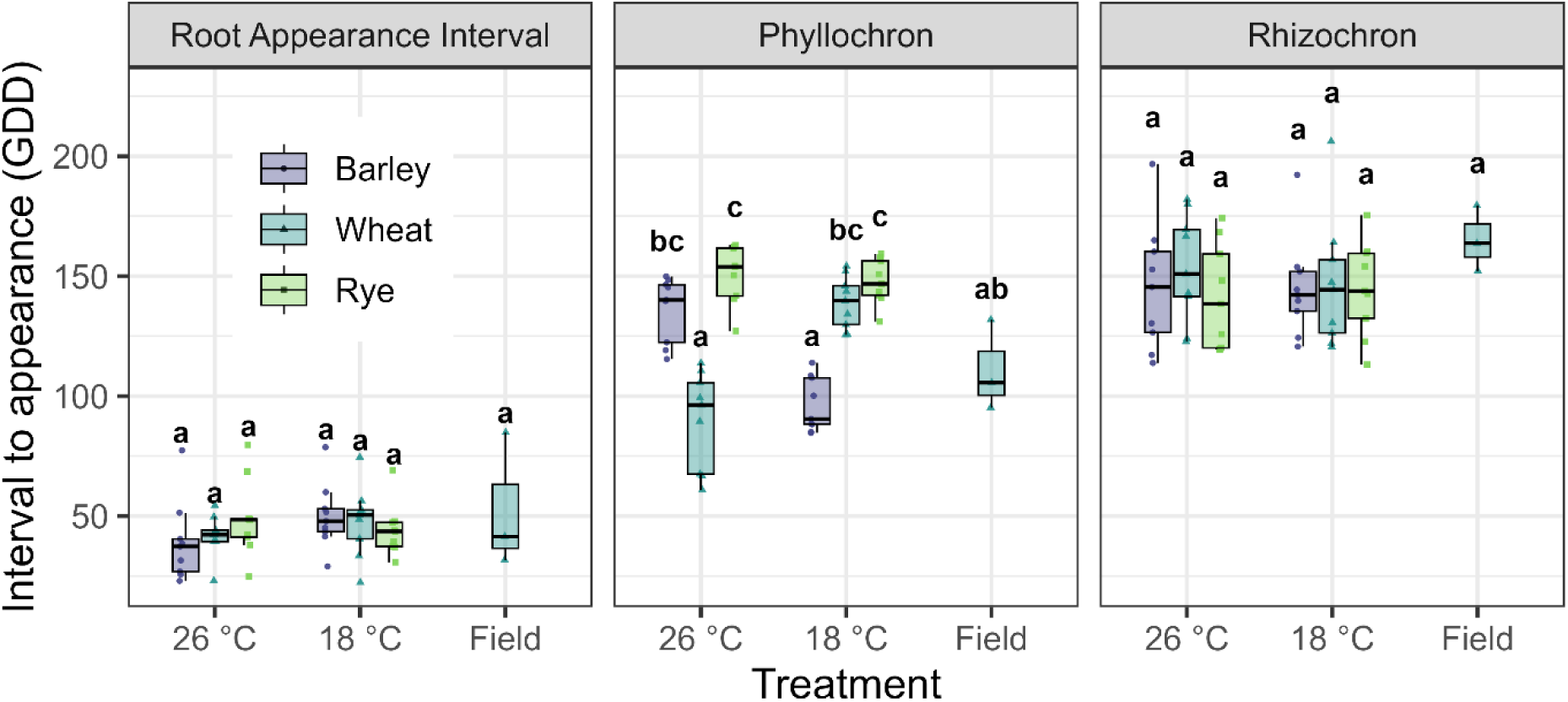
Root Appearance Interval, Rhizochron, and Phyllochron across species and growth conditions. The Root Appearance Interval is the thermal interval requirement (expressed in GDD) between successive root emergence events, irrespective of node. The rhizochron is the thermal interval between emergence of nodal roots from successive stem nodes. The phyllochron is the thermal interval between appearance of successive leaves. In the greenhouse, plants were grown in two temperature conditions 26°C and 18°C. Data from the field (mean crop cycle temperature 15°C) for wheat is also presented. Within each trait, Species × Treatment effects were analyzed using two–way ANOVA (linear model with interaction - lm(Response ∼ Species * Treatment)) followed by Sidak–adjusted pairwise comparisons of estimated marginal means. Different letters within each trait indicate significant differences (P<0.05).

**Table 1.**
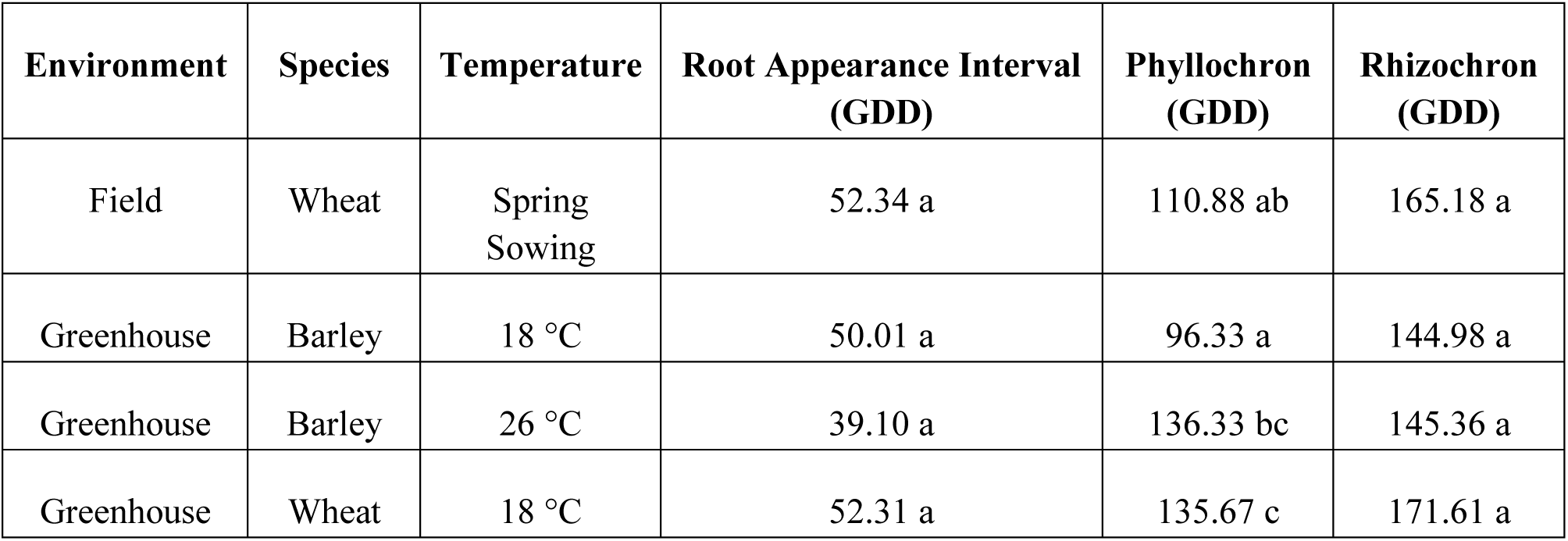
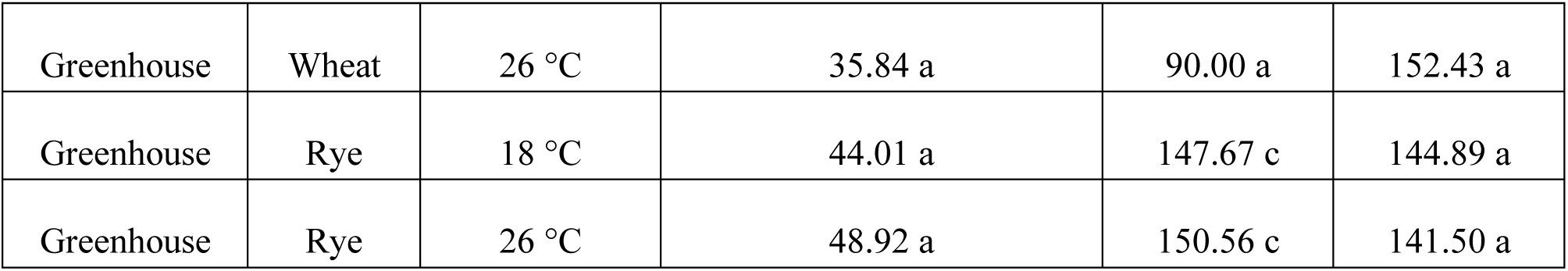
Root and shoot developmental intervals for wheat (*Triticum aestivum* L.), rye (*Secale cereale* L.), and barley (*Hordeum vulgare* L.) measured under field (IPK Gatersleben, Germany) and greenhouse conditions (18 and 26 °C). Values are expressed in thermal time (°C d). The Root Appearance Interval is the thermal interval between successive root emergence events, irrespective of node. The rhizochron is the thermal interval between emergence of nodal roots from successive stem nodes. The phyllochron is the thermal interval between appearance of successive leaves. Different letters within each of these metrics represent significant differences as determined by Tukey’s test (P < 0.05).

| Environment | Species | Temperature | Root Appearance Interval (GDD) | Phyllochron (GDD) | Rhizochron (GDD) |
| --- | --- | --- | --- | --- | --- |
| Field | Wheat | Spring Sowing | 52.34 a | 110.88 ab | 165.18 a |
| Greenhouse | Barley | 18 °C | 50.01 a | 96.33 a | 144.98 a |
| Greenhouse | Barley | 26 °C | 39.10 a | 136.33 bc | 145.36 a |
| Greenhouse | Wheat | 18 °C | 52.31 a | 135.67 c | 171.61 a |
| Greenhouse | Wheat | 26 °C | 35.84 a | 90.00 a | 152.43 a |
| Greenhouse | Rye | 18 °C | 44.01 a | 147.67 c | 144.89 a |
| Greenhouse | Rye | 26 °C | 48.92 a | 150.56 c | 141.50 a |

The thermal requirement for the emergence of roots on a different node consistently exceeded the equivalent thermal requirement for leaf appearance across all observed environments. In the 26 °C greenhouse, the phyllochron ranged from 90.00 to 150.56 °C d leaf^-1^, while rhizochron values were notably higher, ranging from 141.50 to 152.43 °C d node^-1^. In the 18 °C greenhouse, the phyllochron ranged from 86 to 127 °C d leaf^-1^, while rhizochron values were notably higher, ranging from 129.8 to 164.1 °C d node^-1^. This pattern was maintained in the field for spring-sown wheat, where the phyllochron (110.88 °C d) was significantly shorter than the rhizochron (165.18 °C d) (Figure 2, Supplementary Table S1, S3).

In contrast, the root appearance interval, which tracks the emergence of any root axis regardless of its node of origin, was the most rapid developmental metric measured. In the greenhouse, values for this interval ranged from 35.84 to 52.31 °C d root^-1^, averaging 45.3 °C d across all temperature treatments (Figure 2, Supplementary Table S1, S3). The root appearance interval did not show a statistically significant increase at 18 °C compared to 26 °C (p = 0.297), nor were there significant differences between species (p = 0.906) and the species × temperature interaction (p = 0.226) (Supplementary Table S3). Consistent with the stability of both the rhizochron and the root appearance interval, the number of roots per node also remained unaffected by temperature. These data indicate that while the initiation of roots on a specific new stem node (i.e. rhizochron) is a relatively slow process, the plant maintains a high frequency of total root emergence (i.e. root appearance interval) by simultaneously deploying multiple axes across several active nodes. While the phyllochron follows a distinct, species-specific thermal requirement, the root-related metrics (rhizochron and root appearance interval) exhibited higher variability, masking potential treatment effects across the two temperature regimes.

### Root and shoot development diverge in duration and rate

Despite the coordination of shoot and root development with thermal time, a consistent thermal offset (measured in GDD) was observed between the emergence of leaves and nodal roots at the same phytomer. Expressed in thermal time, this offset corresponded to approximately 185.3 °C d between the emergence of a leaf and a root on the same node. Additionally, the trajectories of root and shoot development diverged over the course of the experiment. While leaf appearance on the main stem plateaued at approximately 900 GDD, nodal root emergence on the main stem continued throughout the full 1440 GDD sampling period (Figure 6 and Supplementary Table 2).

The coordination ratio, which quantifies the ratio of the rhizochron and phyllochron, remained relatively stable across a period of 900 GDD. Despite the varying thermal regimes and different species, no significant differences in the coordination ratio were detected (average 0.86) (Table 2). These results indicate that the ratio of organ appearance is highly conserved, maintaining a steady proportion regardless of the specific growth environment or species.

**Table 2.** Ratio between phyllochron and rhizochron (coordination ratio) is presented for each species and temperature condition. The standard deviation (SD) is presented.

| Environment | Species | Temperature | Coordination Ratio | SD |
| --- | --- | --- | --- | --- |
| Field | Wheat | Spring Sowing | 0.68 | 0.14 |
| Greenhouse | Barley | 18 °C | 0.68 | 0.15 |
| Greenhouse | Barley | 26 °C | 0.95 | 0.11 |
| Greenhouse | Rye | 18 °C | 1.04 | 0.17 |
| Greenhouse | Rye | 26 °C | 1.09 | 0.21 |
| Greenhouse | Wheat | 18 °C | 0.97 | 0.18 |
| Greenhouse | Wheat | 26 °C | 0.60 | 0.18 |

### Root developmental timing is stable across temperature regimes

Despite the elevated temperature regime (26 °C), which is generally considered supra-optimal for temperate cereals, both the rhizochron and root appearance interval remained stable. Neither metric exhibited a significant response to temperature (rhizochron: p = 0.839; root appearance interval: p = 0.297), nor were they significantly affected by species. In contrast, the phyllochron exhibited a significant species-by-temperature interaction (p <0.001), indicating that shoot developmental timing was more responsive to thermal variation. These results demonstrate that, unlike shoot development, root developmental timing shows limited sensitivity to temperature variation, maintaining a conserved thermal requirement even under heat stress conditions. This stability was further supported by field observations of wheat, where rhizochron values were consistent with those measured under controlled conditions (Figure 2 and Table 1).

### Root emergence occurs through overlapping activity across multiple nodes

In contrast to leaf appearance, which follows a strict temporal sequence where one leaf emerges before the next begins, root development did not proceed in a strictly sequential manner along the stem axis. Rather than completing root production at one node before initiating the next, plants exhibited overlapping developmental activity across multiple stem nodes. Basal nodes (e.g. stem node 1) continued to produce roots even after the initiation of root emergence at subsequent nodes (e.g. stem nodes 2 and 3) (Figure 3, 4, 5 and 6; Supplementary Table 2). As a result, several nodes contributed simultaneously to total root production throughout development. This concurrent activity provides a mechanistic explanation for the relatively short root appearance interval despite the longer rhizochron, as multiple nodes are actively producing roots at any given time. These findings indicate that in temperate cereals, although phytomers are initiated sequentially, their subsequent root emergence proceeds in a coordinated and overlapping manner rather than as a strictly discrete, node-by-node progression.

**Figure 3.**
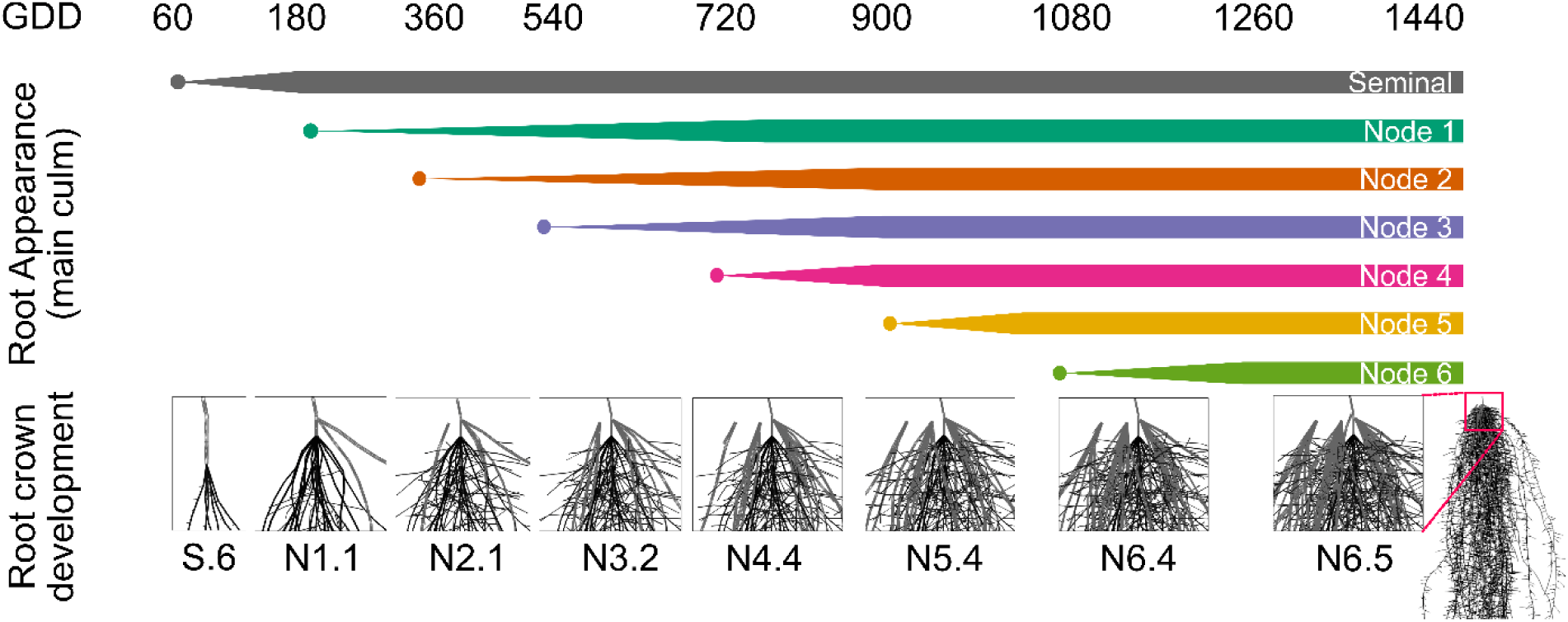
The appearance of seminal and nodal roots from different stem nodes on the main stem. The rhizochron, the appearance of roots on successive stem nodes, is linked to thermal time. The development of the root crown (enlarged image of the root crown simulated with *OpenSimRoot*) showing the development of roots on the main stem. Root developmental stages were classified using a root scale in which the first number represents the number of main stem nodes with emerged roots, and the second number indicates the number of nodal roots on the most recently developed node. The onset of root appearance at each node is indicated by colored dots, while the increase in root production is represented by progressively widening triangles, reaching a right-angled form at maximum root number.

**Figure 4.**
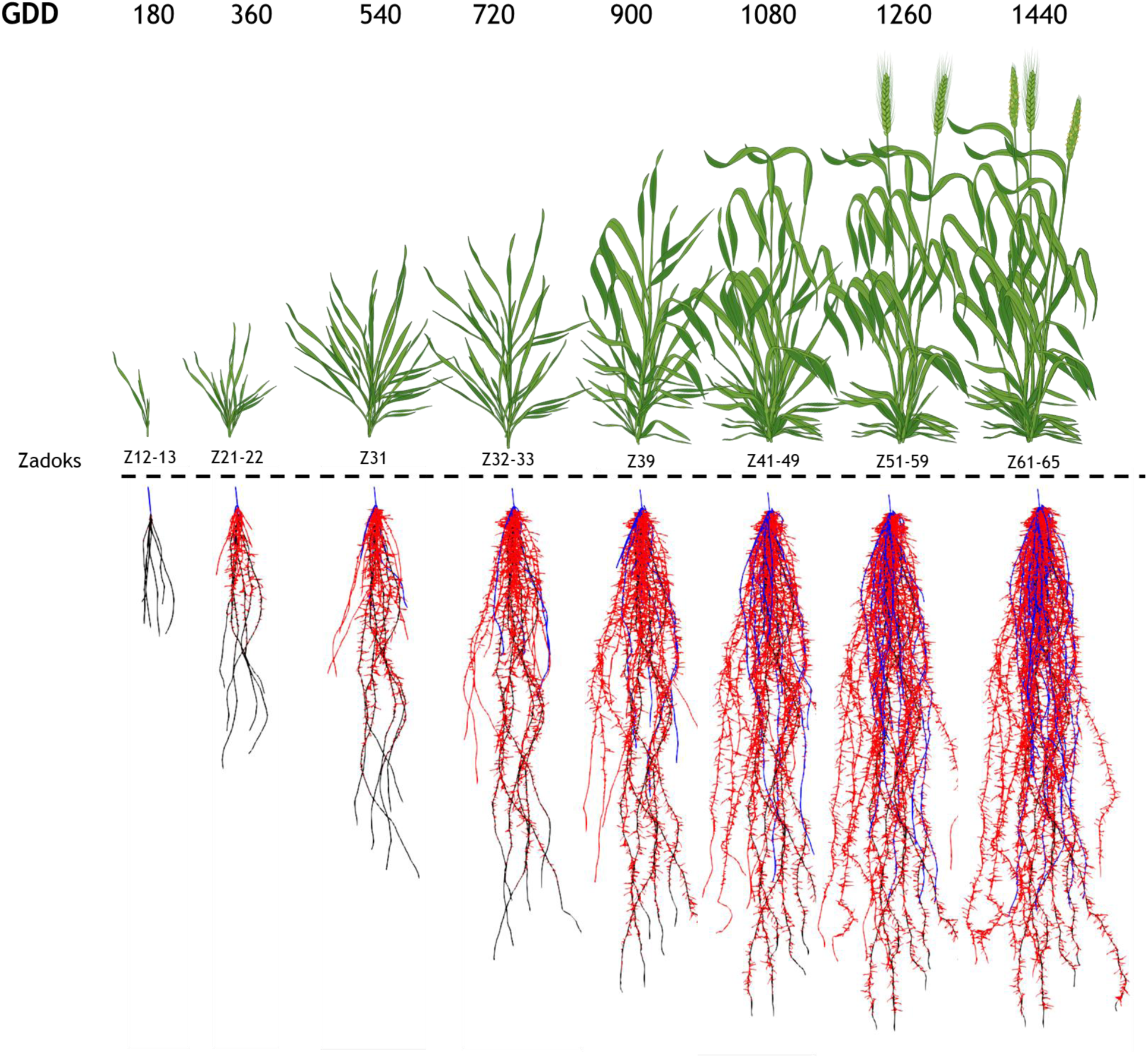
Above-ground development stages according to the Zadoks scale and corresponding below-ground development simulated with *OpenSimRoot* (Postma et al. 2017). The scheme summarizes the mean observations from the two temperature regimes evaluated (18 °C and 26 °C in the greenhouse), considering the three species and the three genotypes per species across eight sampling dates (180-1440 GDD) (N=432). Seminal roots (brown), nodal roots on the main stem (red), nodal roots on tillers (blue). Schematic representations of the Zadoks scale were generated using BioRender.

**Figure 5.**
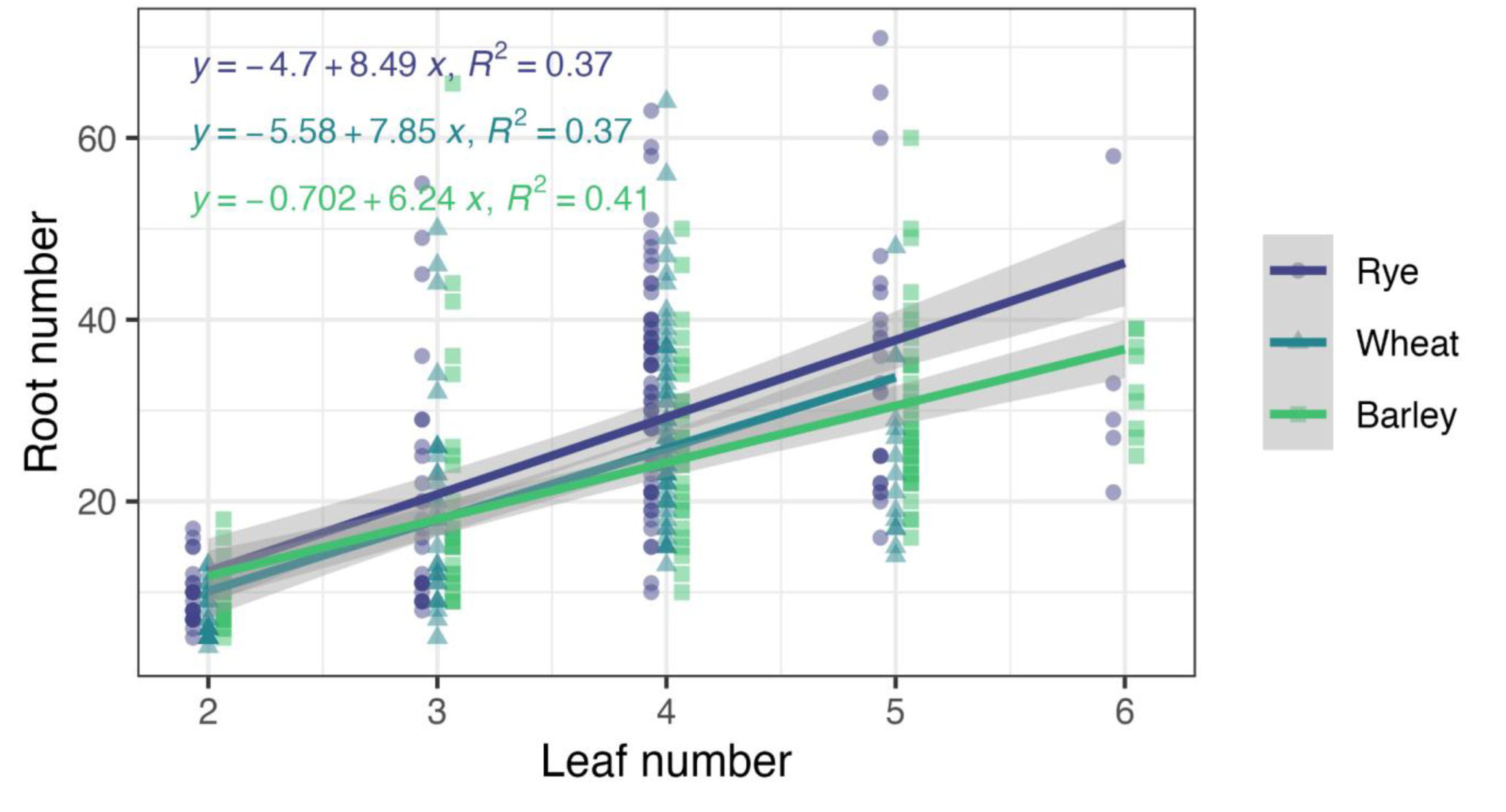
Regression of root and shoot number on the main stem. For each species, values represent the mean of the two environments evaluated (18 °C and 26 °C). Data points include all sampling times (N=432). Linear model using ordinary least squares was fitted per species and shaded band represents 95% CI.

**Figure 6.**
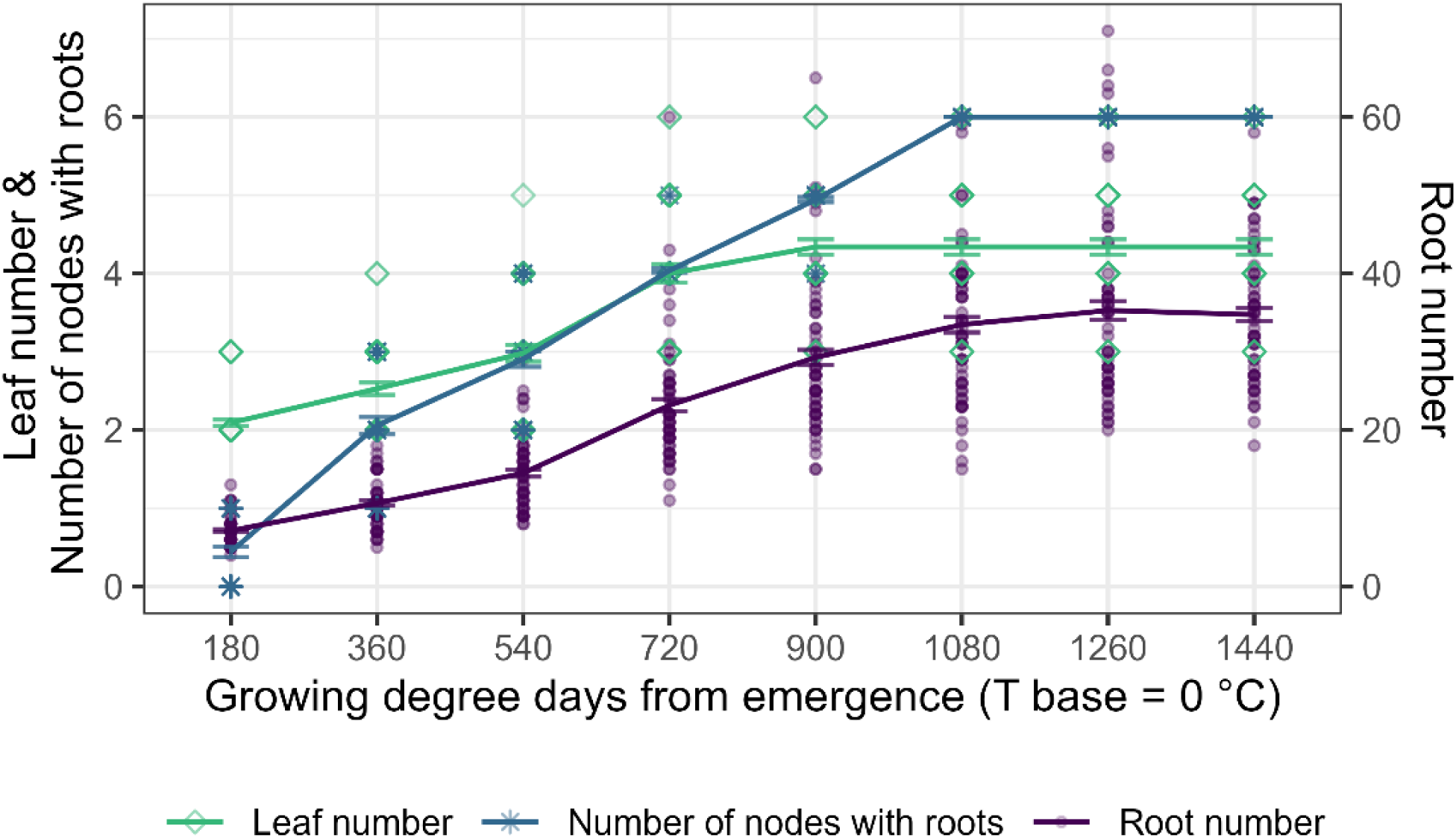
Dynamics of leaf number, root number, and number of nodes with roots on the main stem across the crop cycle. Lines represent the mean of the two environments evaluated (18 °C and 26 °C in the greenhouse), considering the three species and the three genotypes per species across eight sampling dates (180-1440 GDD) (N=432). Error bars represent the standard error.

The rate of root production within individual nodes varied systematically with node position along the stem. Proximal nodes (e.g. stem node 1) were active in root emergence over a longer thermal interval (GDD) i.e. required a greater accumulation of GDD to reach their final occupancy, compared to more distal stem nodes (Figure 3/Table S2). However, each stem node had a similar final root occupancy i.e. number of roots per node. This spatial and temporal variation indicates that root production dynamics are not uniform across phytomers, but instead depend on node position. These differences suggest that developmental constraints or regulatory factors vary along the stem axis, potentially reflecting gradients in tissue maturity, assimilate availability, or hormonal signaling.

## Discussion

### A conserved, hierarchical framework for root system development

This study establishes that root development in temperate cereals follows a conserved thermal program that is tightly coordinated with shoot phenology through the phytomer. Although the phytomer concept has long suggested a coordinated development of roots and shoots in cereals and forage grasses, previous studies have only partially characterized root developmental dynamics and lack a quantitative framework across species equivalent to that used for shoot phenology (Porter et al., 1986; Vincent and Gregory 1989 a, b; Klepper et al., 1997; Asseng et al. 1997; Nemoto et al. 1995; Yang et al. 1998; Watt et al. 2009; Singh et al. 2010; Robin et al 2016; Robin et al 2021). By defining the rhizochron as the thermal interval between the emergence of roots from successive nodes, we provide a quantitative descriptor of below-ground development that parallels the phyllochron. Together with the root appearance interval, these metrics reveal a hierarchical developmental system in which node activation and root axis production operate at distinct but coordinated developmental scales.

### Overlapping phytomer activity drives continuous root system expansion

Root emergence was not strictly sequential but instead characterized by overlapping developmental activity across multiple phytomer positions. Early-initiated phytomers continued producing roots after subsequent nodes were activated, resulting in simultaneous contributions to the total root system in acropetal sequence. This challenges the traditional stepwise view of root development and supports a model in which expansion is driven by coordinated, concurrent phytomer activity, similar to perennial ryegrass and tall fescue (Matthew et al 2001; Robin et al 2021).

### Node position shapes root production dynamics

In addition to temporal coordination, root development exhibited clear spatial differences along the stem axis. The rate of root production depended on node position. Proximal nodes (e.g., node 1) remained active longer and required greater GDD to reach final occupancy, whereas more distal nodes produced roots more rapidly. This may reflect increasing resource availability as shoot biomass and photosynthetic capacity rise during development (Schierenbeck et al. 2016; Slafer et al., 2022), or increasing mechanical demands of supporting a larger shoot.

The timing of adventitious root emergence may be further modulated by the anatomical maturity of the node. For example, in rice, in more mature nodes like the third or fourth nodes, emergence occurs more rapidly because root primordia tips are often already in contact with the epidermis. In younger apical nodes (e.g. node 6), the primordia remain sequestered beneath two to four layers of parenchymal cells, which delays emergence. These adventitious roots represent a critical physiological adaptation to environmental stress. By facilitating gas exchange, they enable the plant to mitigate the oxygen deficiencies characteristic of submergence and flooding (Steffens et al. 2012). Regardless of the underlying driver, these results underscore that a single parameter cannot adequately describe root system architecture. Capturing the full dynamics of root formation requires accounting for both the timing of nodal activation and the intrinsic rate of production within those nodes.

Notably, this specialization extends to morphology and function. While temperate cereals employ more uniform nodal occupancy than maize (York and Lynch, 2015), anatomy and stress responses vary by position (York and Lynch, 2015; Yang et al 2019; Koehler et al 2025). These distinctions likely relate to variations in nutrient uptake, AMF colonization, or disease resistance (Lynch, 2019). Similar gradients exist in the canopy: distal phytomers, particularly those associated with the flag leaf, possess larger blades and contribute disproportionately to photosynthesis (Evans, 1983; Reynolds et al., 2012a,b; Schierenbeck et al. 2024; Ruz-Ruiz et al., 2026), accompanied by stratification in nitrogen content (Dreccer et al., 2000). Ultimately, the coordination of distinct anatomical and morphological traits across different phytomer positions reflects a sophisticated functional mapping, allowing the plant to meet the shifting demands of its life cycle.

### Phenological coordination of the phytomer

The lack of significant variation in the coordination ratio across species and temperatures suggests that the timing of leaf and nodal root emergence is governed by a strictly coupled developmental program. While absolute rates of growth may accelerate with temperature, the coordination ratio appears to be a fixed trait in annual temperate cereals, however may change over seasons in perennial cereals (Robins et al 2021). This constancy implies that for every unit of shoot development, a proportional and predictable amount of root development occurs, regardless of whether the plant is experiencing optimal (18 °C) or supra-optimal (26 °C) temperatures affecting the above- and below-ground growth rate across the crop cycle (Langridge & Reynolds, 2021) (Supplementary Figure S2).

Within this strictly coupled program, a consistent developmental offset dictates the temporal hierarchy of individual phytomers. Calculated against the mean greenhouse phyllochron, our observed developmental delay of 185 GDD equates to a lag of approximately 1.5 leaves; this means root emergence at a given node begins only after 1.5 subsequent leaves have developed. While this value is lower than the two to four leaf delay previously reported for initial nodal root emergence (Fujii, 1961; Vincent and Gregory, 1989 a, b; Rebouillat et al., 2009; Watt et al., 2009; Singh et al., 2010; Robin et al. 2021), the fundamental developmental logic remains consistent: the leaf maintains a significant developmental head start over the roots of the same phytomer. A structurally homologous offset is well documented in the above-ground architecture of grasses concerning tiller emergence. For instance, Evers et al. (2005) characterized a similar ‘tiller-appearance delay’ between the appearance of leaves and their corresponding axillary tillers. This parallel suggests a generalized developmental rule in Poaceae: the emergence of lateral organs, whether nodal roots for resource acquisition or tillers for structural expansion, is subject to a highly conserved thermal delay following the appearance of the primary photosynthetic organ at that node.

The greater thermal requirement of the rhizochron, which necessitates this delay, likely reflects fundamental differences in organogenesis between above- and below-ground structures. In temperate cereals, a set number of leaf primordia are pre-formed within the embryo (Ochavagia et al., 2018), meaning their subsequent appearance is primarily a function of cell elongation. In contrast, nodal roots are not pre-formed; they must be newly initiated from stem nodes before they can emerge (Kersetter & Hake, 1997).

Interestingly, while the rates of root and shoot development are coordinated through these conserved thermal intervals early on, their durations diverge as accumulated thermal time progresses. Leaf appearance on the main stem ceased at approximately 900 GDD, whereas nodal root axis emergence continued until the booting stage (1080 GDD). This late-stage phenological decoupling indicates that the developmental program governing root emergence persists beyond the completion of leaf emergence on the main stem.

### Axial root emergence is not affected by thermal stress

A critical factor in interpreting these results is the disparity between shoot and root thermal environments (Jamieson et al., 1998). Under field conditions, soil temperature is typically more buffered and often lower than air temperature (Ashcroft & Gollan, 2013). However, we observed consistent rhizochron and root appearance interval values across both controlled and spring field systems. This consistency suggests that the timing of nodal root emergence is regulated at the shoot apical meristem or stem nodes, where temperatures track ambient air conditions more closely than the bulk soil. It is well-established that above-ground development is governed by thermal time sensed at the shoot apical meristem (Jamieson et al., 1995; Jamieson et al., 1998; McMaster et al., 2003). Because the vegetative apex remains near the soil surface until stem elongation, it is uniquely positioned to integrate the temperature of the crown environment. Our findings suggest that this same sensing mechanism dictates the cadence of nodal root emergence. It is important to distinguish this developmental timing from the growth or elongation of the root itself; while the timing of emergence appears linked to the shoot apical meristem, the subsequent growth and elongation of the root are known to be directly sensitive to local temperature conditions (Ai et al 2023).This model is further supported by the fact that a thermal time-scale using air or soil surface temperatures adequately describes nodal root axis production across diverse sowing dates and sites, whereas incorporating deeper soil temperatures provides no significant improvement in predictive accuracy (Vincent and Gregory, 1989a, b).

The stability of these metrics under elevated temperature further supports a conserved developmental program. Unlike the more plastic phyllochron, root developmental timing showed limited responsiveness to temperature, indicating partial decoupling in sensitivity between root and shoot development. Environmental stress may still modulate expression; for example, root emergence can be reduced under drought (Yamauchi et al., 1994).

Together, these findings support a model in which the regulation of nodal root emergence is governed by systemic developmental signals linked to shoot phenology rather than local soil conditions. Because nodal roots originate from stem nodes, their timing is likely coordinated with the same genetic pathways controlling above-ground development. In temperate cereals, these include vernalization (VRN), photoperiod (PPD), and earliness per se (EPS) loci in wheat (Distelfeld et al., 2009; Ochagavía et al., 2018; Dixon et al., 2018), with homologous systems in barley (Turner et al., 2005; Digel et al., 2015) and rye (Rabanus-Wallace et al. 2021). Specifically, regulators like *ELF3* maintain photoperiod sensitivity by suppressing both *FT1* expression and the production of bioactive gibberellins (GA), which are essential for coordinating inflorescence development with vegetative growth. Furthermore, root architecture is fine-tuned by dosage-sensitive pathways, such as the *OPRIII* gene cluster, which modulates seminal and nodal root elongation through the biosynthesis of jasmonic acid (JA-Ile) and the regulation of reactive oxygen species (ROS) distribution (Boden et al 2014; Gabay et al 2023). Accordingly, the phyllochron, rhizochron, and root appearance interval may be interpreted as integrative traits emerging from shared genetic control of phenology.

Because nodal roots originate from stem nodes, their emergence is governed by the same genetic pathways that regulate shoot phenology. Transcription factors such as ARF7 and ARF19 serve as cellular translators of these thermal intervals, bridging the gap between temperature signals and physical growth (Borniego et al., 2026). These proteins undergo rapid and reversible partitioning into the nucleus when temperatures rise, a mechanism that directly promotes the cell elongation necessary for nodal root axes to emerge from the stem. The requirement to move from de novo initiation through to physical emergence may introduce a systematic delay, while the molecular activity provides a concrete cellular basis for how temperate cereals maintain a unified developmental pace.

Although the experiments were conducted under controlled conditions, the agreement with field observations reinforces the robustness of this thermal framework. Nevertheless, variation in rhizochron and root appearance interval is likely influenced by genotypic differences in phenological regulation. Further work is needed to assess the stability of these relationships across genotypes with contrasting developmental strategies (e.g. winter versus spring types, photoperiod sensitivity) and under a wider range of environmental conditions.

### Implications for modeling and breeding

The identification of the rhizochron provides a foundation for incorporating root development into functional-structural plant models such as *OpenSimRoot* (Postma et al., 2017). Currently, versions of OpenSimRoot simulate general plant growth based on thermal time (Schäfer, 2022); however, they often lack an explicit, stage-based framework for root development specifically. By representing root system development and expansion as a sequence of thermally regulated developmental events, this framework enables models to move beyond static representations based on biomass or root length (Barraclough, 1984; Barraclough and Leigh, 1984; Porter et al., 1986; Klepper et al., 1997) toward dynamic, stage-based growth processes. This approach also offers opportunities to simplify model parameterization. Rather than treating root and shoot development as independent processes, a shared thermal framework allows coordinated simulation of organ emergence based on common developmental timing.

Beyond computational modeling, integrating the rhizochron into crop improvement strategies provides a much-needed phenological context for trait-based breeding. While root system establishment begins early (180–540 GDD), the physiological priority shifts during the critical yield-determination phase (720–1440 GDD), where grain number per m^2^ is established (Slafer & Savin, 1994; Satorre & Slafer, 1999; Carrera et al. 2024). Because late-emerging nodal roots require significant carbon allocation precisely when the developing spike is the dominant sink, their contribution to late-season resource acquisition must be carefully weighed against their metabolic cost. The primary breeding objective is to prioritize genotypes with resource-efficient traits, such as reduced metabolic costs through cortical aerenchyma, root cortical senescence, and optimized spatial distribution through steep nodal angles (Schneider et al. 2017, 2022, 2023), to support the high demands of stem elongation and flowering without creating a carbon sink that competes with seed filling. Identifying genotypes that maximize uptake efficiency through anatomical and structural resilience remains a primary challenge for unifying above- and below-ground phenological frameworks. By integrating the rhizochron and root appearance interval into this model, we can more precisely tailor these optimal anatomical and architectural traits to specific plant growth stages, ensuring that root function is synchronized with the evolving resource demands and stress profiles of the crop’s life cycle.

Extending thermal-time-based phenological frameworks below ground offers a pathway to unify developmental descriptions across crop species. While above-ground phenology is well characterized in many crops such as maize, rice, and soybean (Ritchie & Hanway 1982; Counce et al., 2000; Fehr & Caviness, 1977), equivalent descriptors for root development remain limited. The extent to which the rhizochron concept applies to species with contrasting architectures, such as taprooted dicots or low-tillering cereals, remains an important direction for future research.

## Conclusions

Defining the rhizochron establishes a quantitative framework for root development that complements shoot phenology. Integrating this metric into whole-plant models has the potential to enable a unified view of coordinated above- and below-ground development, supporting improved crop modeling and breeding strategies.

## Author Contributions

MS and HMS conceived and designed the study. Plant growth and wet-lab were performed by MS, AT, ILV, DHJ, PL, AW and CT. Solution culture set-up was designed by MS, HMS, ILV and DJ. Dry-lab methodology was performed by MS, PL, AW and CT. MS, HMS, AT and ILV performed data analyses. Funding acquired by HMS. MS and HMS drafted the manuscript. All authors contributed substantially to the revisions.

## Funding

This study was kindly supported by the Leibniz Association (Resilient Roots to HMS and MS). Open Access funding enabled and organized by Projekt DEAL.

Declarations

## Data availability

Data available in article supplementary material

## Acknowledgements

We thank Prof. Dr. Andreas Boerner from the German Ex-Situ Genebank (IPK Gatersleben, Germany) for providing the genetic resources. We thank Luisa Meier, Luisa Wolfgramm, Maria Schoen, Rashmi Aryal, Osalumen Philemon Irusota, Wirassaya Tobdan, Gopika Shaju, Misbah Chaudhry and Mohsen Morovati for their assistance with plant evaluations. We thank Peter Schreiber, Enk Geyer, Kerstin Jacobs, Anke Mueller and Kathrin Tiemann for their assistance with greenhouse experiment management.

