## Supplemental Information for "A thermal time framework drives coordinated below- and above-ground development in temperate cereal crops"

**Supplementary information**

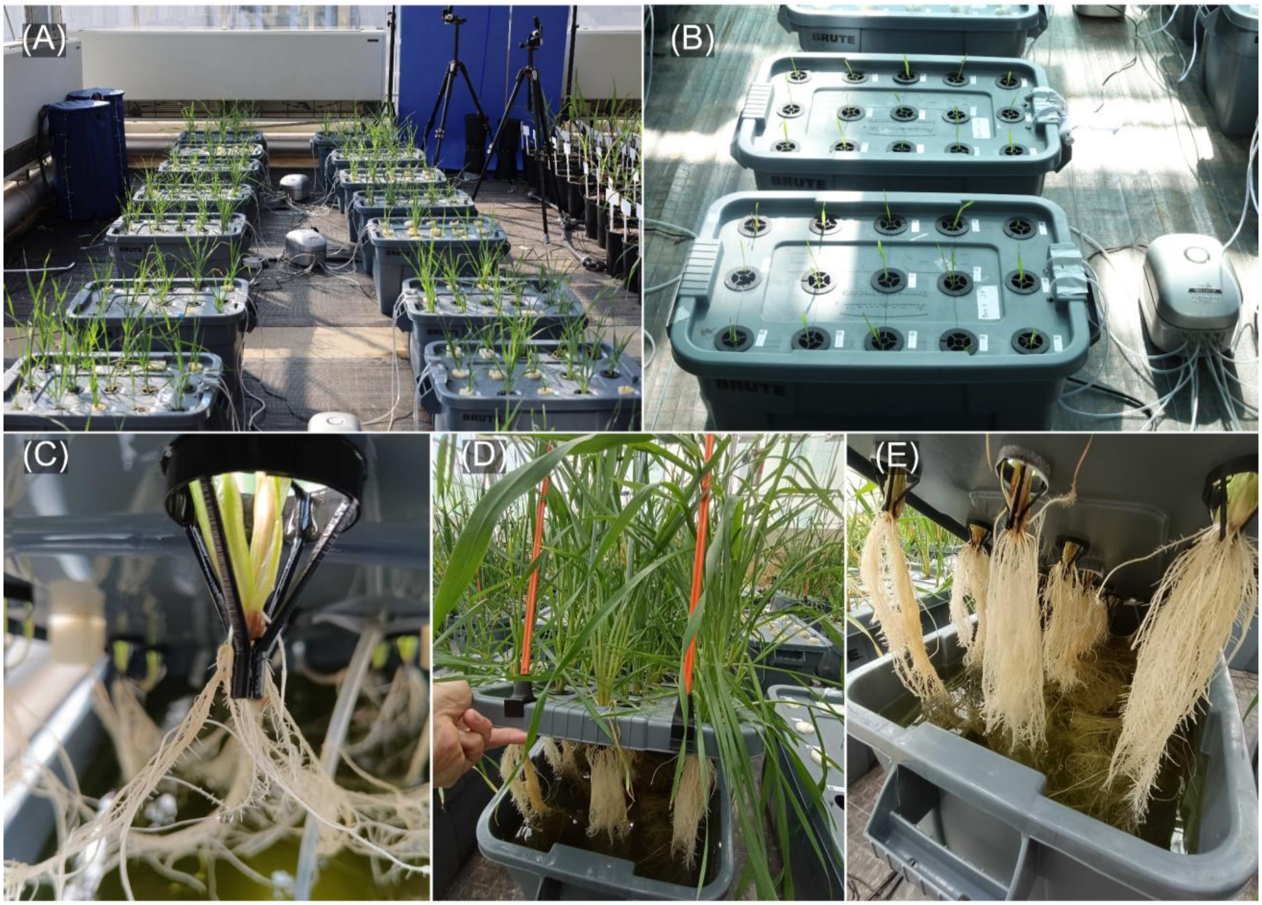


**Figure S1.** Hydroponics experiment setup. The photos represent the deep-water-culture hydroponic setup with containers and aerators used during the experiment (A, B) and plants were held in place using a plastic holder (C) while roots growing in a nutrient solution (D, E).

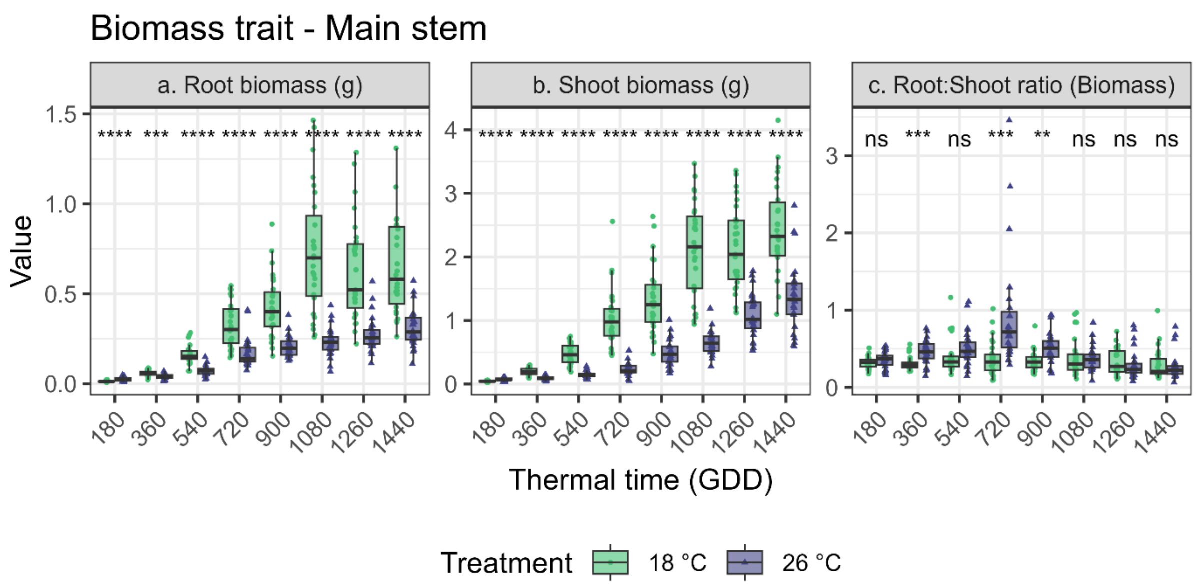


**Figure S2.** The data points represent the dynamics of root biomass (a), shoot biomass accumulation (b), and root to shoot biomass ratio through 8 sampling times (every 180 GDD) of 9 genotypes of spring wheat, barley and rye under two temperature conditions. * indicate significance P<0.05; **P<0.01; ***P<0.001; ****P<0.0001; ns (no significance) when both temperature regimes were compared at a given sampling time (N=54).

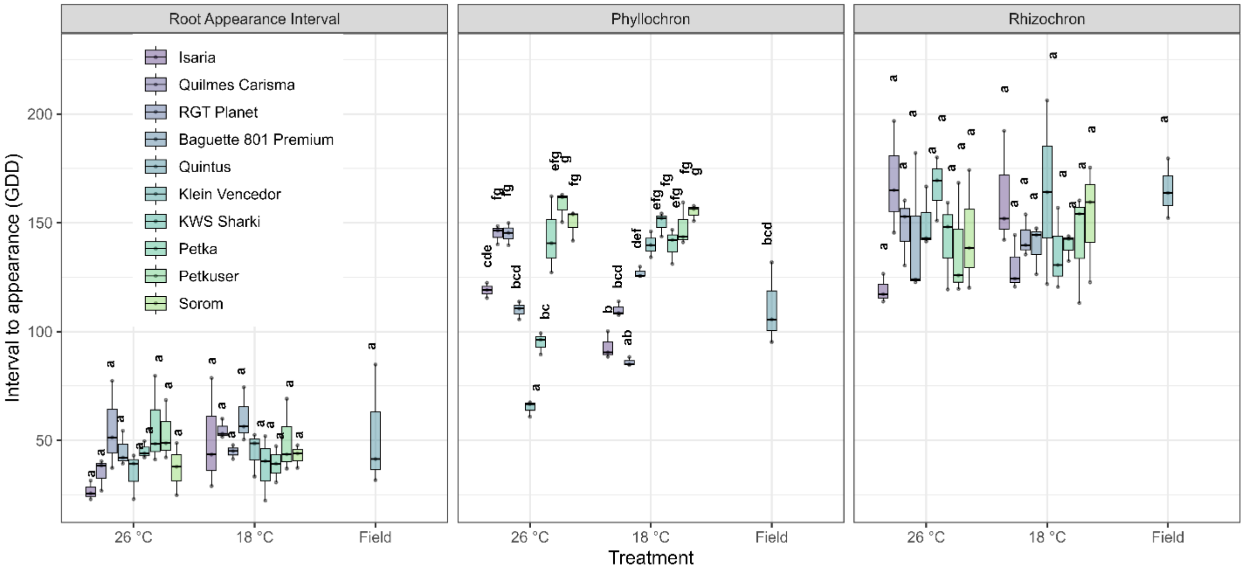


**Figure S3.** Mean Root Appearance Interval, , Phyllochron and Rhizochron across genotypes and growth conditions. In the greenhouse, plants were grown in two temperature conditions 26 °C and 18 °C. Data from the field (mean crop cycle temperature 15 °C) for wheat is also presented. There are three barley (Isaria, Quilmes Carisma, RGT Planet), four wheat (Baguette 801 Premium, Quintus, Klein Vencedor, KWS Sharki) and three rye (Petka, Petkuser, Sorom) genotypes. Within each trait, Genotype × Treatment effects were analyzed using two-way ANOVA [linear model with interaction - lm(Response ~ Genotype × Treatment)] followed by Sidak-adjusted pairwise comparisons of estimated marginal means. Different letters within each trait indicate significant differences ($P<0.05$).

**Table S1.** Root and shoot developmental intervals for wheat, rye, and barley measured in greenhouse conditions under two temperature regimes (18 and 26 °C). Values are expressed in thermal time (°C d). The Root Appearance Interval is the thermal interval between successive root emergence events, irrespective of node. The rhizochron is the thermal interval between the emergence of nodal roots from successive stem nodes. The phyllochron is the thermal interval between the appearance of successive leaves. Different letters within each metric indicate significant differences, as determined by Tukey's test ($P < 0.05$).

| **Environment** | **Species** | **Temperature** | **Genotype** | **Root Appearance Interval** | **Phyllochron** | **Rhizochron** |
|---|---|---|---|---|---|---|
| Field | Wheat | Spring | Quintus | 52.34 a | 110.88 def | 165.18 a |
| Greenhouse | Barley | 18 °C | Isaria | 50.39 a | 93.00 fg | 162.12 a |
| Greenhouse | Barley | 18 °C | Quilmes Carisma | 54.86 a | 110.00 defg | 129.80 a |
| Greenhouse | Barley | 18 °C | RGT Planet | 44.78 a | 86.00 gh | 143.02 a |
| Greenhouse | Barley | 26 °C | Isaria | 26.70 a | 119.00 cde | 119.18 a |
| Greenhouse | Barley | 26 °C | Quilmes Carisma | 35.24 a | 145.00 ab | 169.09 a |
| Greenhouse | Barley | 26 °C | RGT Planet | 55.36 a | 145.00 ab | 147.82 a |
| Greenhouse | Wheat | 18 °C | Baguette 801 | 60.39 a | 127.00 bcd | 139.38 a |
| Greenhouse | Wheat | 18 °C | Klein Vencedor | 44.84 a | 140.00 abc | 164.07 a |
| Greenhouse | Wheat | 18 °C | Sharki | 38.24 a | 150.00 ab | 136.00 a |
| Greenhouse | Wheat | 26 °C | Baguette 801 | 45.27 a | 110.00 defg | 142.93 a |
| Greenhouse | Wheat | 26 °C | Klein Vencedor | 35.12 a | 65.00 h | 150.37 a |
| Greenhouse | Wheat | 26 °C | Sharki | 45.31 a | 95.00 efg | 166.86 a |
| Greenhouse | Rye | 18 °C | Petka | 39.10 a | 140.00 abc | 139.62 a |
| Greenhouse | Rye | 18 °C | Petkuser | 49.90 a | 148.00 ab | 142.50 a |
| Greenhouse | Rye | 18 °C | Sorom | 43.01 a | 155.00 a | 152.54 a |
| Greenhouse | Rye | 26 °C | Petka | 56.45 a | 143.34 ab | 142.27 a |
| Greenhouse | Rye | 26 °C | Petkuser | 53.15 a | 158.33 a | 137.95 a |
| Greenhouse | Rye | 26 °C | Sorom | 37.16 a | 150.00 ab | 144.27 a |

**Table S2.** *OpenSimRoot* Parameters. This table represents the average values of the number of roots per node, nodal root appearance on each node, and the total root length across species and greenhouse conditions. These parameters were used in the functional-structural plant model *OpenSimRoot* to simulate root growth and development at eight points according to cumulative GDD.

| **Time (days)** | **8** | **16** | **24** | **32** | **40** | **48** | **56** | **64** |
|---|---|---|---|---|---|---|---|---|
| **Total root length (cm)** | 412.61 | 1508.82 | 3148.64 | 7242.38 | 11570.19 | 14262.47 | 15170.01 | 19585.08 |
| **GDD** | 180 | 360 | 540 | 720 | 900 | 1080 | 1260 | 1440 |
| **Zadoks** | Z12-13 | Z21-22 | Z31 | Z32-33 | Z39 | Z45 | Z55 | Z65 |
| **Seminal** | 6 | 6 | 6 | 6 | 6 | 6 | 6 | 6 |
| **Node 1** | 1 | 2 | 3 | 4 | 4 | 4 | 4 | 4 |
| **Node 2** | 0 | 1 | 2 | 4 | 5 | 5 | 5 | 5 |
| **Node 3** | 0 | 0 | 2 | 4 | 5 | 5 | 5 | 5 |
| **Node 4** | 0 | 0 | 0 | 4 | 5 | 5 | 5 | 5 |
| **Node 5** | 0 | 0 | 0 | 0 | 4 | 5 | 5 | 5 |
| **Node 6** | 0 | 0 | 0 | 0 | 0 | 4 | 5 | 5 |
| **Tillers** | 0 | 2 | 4 | 10 | 23 | 38 | 48 | 52 |

**Table S3.** Summary of probabilities (p-values) from a Two-Way Analysis of Variance (ANOVA) evaluating the effects of Species (Barley, Rye, Wheat), Temperature (18 °C, 26 °C), and their Interaction (Species x Temperature) on three cereal growth metrics: Root Appearance Interval, Phyllochron, and Rhizochron. Data represents greenhouse-grown plants (n = 9 replicates per species).

| **Source of Variation** | **Root Appearance Interval** (p-values) | **Phyllochron** (p=values) | **Rhizochron** (p-values) |
|---|---|---|---|
| **Species** | 0.906 | < 0.001 | 0.674 |
| **Temperature** | 0.297 | 0.578 | 0.839 |
| **Species × Temp** | 0.226 | < 0.001 | 0.800 |
